# Genome size estimation from long read overlaps

**DOI:** 10.1101/2024.11.27.625777

**Authors:** Michael B Hall, Lachlan J M Coin

## Abstract

**Summary:** Accurate genome size estimation is an important component of genomic analyses, though existing tools are primarily optimised for short-read data. We present LRGE, a novel tool that uses read-to-read overlap information to estimate genome size in a reference-free manner. LRGE calculates per-read genome size estimates by analysing the expected number of overlaps for each read, considering read lengths and a minimum overlap threshold. The final size is taken as the median of these estimates, ensuring robustness to outliers such as reads with no overlaps. Additionally, LRGE provides an expected confidence range for the estimate. LRGE outperforms *k*-mer-based methods in both accuracy and computational efficiency and produces genome size estimates comparable to those from assembly-based approaches, like Raven, while using significantly less computational resources. We validate LRGE on a large, diverse bacterial dataset and confirm it generalises to eukaryotic datasets.

**Availability and implementation:** Our method, LRGE (**L**ong **R**ead-based **G**enome size **E**stimation from overlaps), is implemented in Rust and is available as a precompiled binary for most architectures, a Bioconda package, a prebuilt container image, and a crates.io package as a binary (`lrge`) or library (`liblrge`). The source code is available at https://github.com/mbhall88/lrge under an MIT license.

## Introduction

Genome size is a fundamental genomic characteristic essential for downstream analyses such as genome assembly (1) and evolutionary studies (2), its accurate estimation remains challenging, especially for non-model organisms and datasets with high heterogeneity or repetitive content. Existing methods primarily focus on short-read data and often require high computational resources or rely on pre-assembled references (3; 4), limiting their applicability to modern long-read sequencing platforms from Pacific Biosciences (PacBio) and Oxford Nanopore Technologies (ONT).

The advancement of long-read sequencing technologies has made it relatively easy to generate high-quality bacterial genome assemblies (5). Combined with the ever-increasing throughput of sequencing, automated pipelines for tasks like variant calling and genome assembly are now common (6; 7; 8). These pipelines often require a genome size estimation from the user (6), or optionally estimate one (7; 8). However, existing size estimation tools are typically designed for short reads (4; 3) and struggle with the higher error rates of long reads, which can introduce many erroneous *k*-mers, the basis for some estimation methods (9; 4). Additionally, determining sequencing depth requires knowledge of the genome size (10), so pipelines that downsample sequencing data to improve computational efficiency must either rely on user-provided size estimates or calculate them automatically.

Here, we present a novel approach leveraging long-read overlap data to provide accurate genome size estimates in a reference-free manner, without a reliance on *k*-mers. By focusing on read-to-read overlaps, our method efficiently captures genome-wide patterns of coverage and redundancy, offering a robust alternative to *k*-mer-based and assembly-dependent techniques. This tool is designed for analysing long-read datasets, making genome size estimation accessible for a broader range of organisms and experimental contexts.

Our method, LRGE, estimates genome size by analysing how individual reads overlap with one another. For each query read, it calculates the expected number of overlaps with a set of target reads, considering both the lengths of the reads and a minimum overlap threshold. Using this overlap information, the method derives a genome size estimate for each query read. These per-read estimates are then aggregated using the median, ensuring robustness to outliers such as reads with no overlaps, which would otherwise produce infinite estimates. This approach leverages the statistical properties of read overlaps to infer genome size without requiring a pre-assembled reference.

## Methods

Suppose the genome size is **GS**. We assume that we have sequenced a set of |*T*| target reads *T* with average length 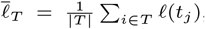, where *ℓ*(*t*_*j*_) is the length of a specific read *t*_*j*_ ∈ *T*; as well as a second set of |*Q*| query reads *Q* with average read length 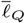, defined in the same manner as 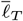. We denote by **ov**(*Q, T*) the set of reads in *Q* which overlap with *T* and by *T* \ {*q*_*i*_} the set of reads in *T* not including *q*_*i*_, which enables us to consider the case that *Q* and *T* may include the same reads.

The probability that a read *q*_*i*_ overlaps with a distinct read *t*_*j*_ is equal to

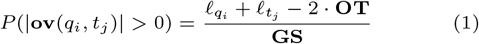

where **OT** is the base-pair threshold on the minimal overlap length, as defined by the minimap2 minimum chaining score, −m, (default 100bp for overlaps) (11). The expected count of overlaps is thus given by

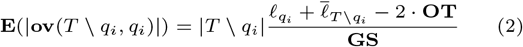

which gives us a per-read estimate of **GS**,

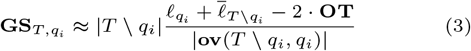

We calculate the overall genome size estimate, **GS**_*T,Q*_, as the **median** of the finite per-read estimates, 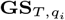, for all reads *q*_*i*_ ∈ *Q*

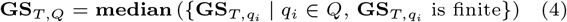

as a read *q*_*i*_ with no overlaps will return a per-read estimate of ∞.

### Implementation

We have implemented this estimation process in the software tool LRGE using the Rust programming language, with overlaps being generated by minimap2 (11). LRGE offers a command-line application, along with a library API for use in other Rust projects.

The implementation provides two strategies for estimation: Two-set (*2set*), whereby *Q* and *T* are disjoint sets of reads; and all-vs-all (*ava*), where *Q* and *T* are identical sets of reads.

The *2set* strategy has the advantage where *Q* can be made smaller than *T* to reduce the number of per-read estimates, 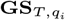, as we ultimately take the median of these estimates. This strategy is the default used in LRGE, with |*Q*| = 5000 and |*T*| = 10000.

In the *ava* strategy, we ignore self-overlaps. Both strategies count only the first overlap between each pair of reads, ignoring subsequent overlaps.

### Evaluation

We compared the genome size estimations from LRGE using the *ava* and *2set* strategies, as well as GenomeScope2 (4), Mash (9), and Raven (12).

For LRGE *ava* we used 25,000 randomly selected reads, while for the *2set* strategy, we used a target set size of 10,000 and a query set size of 5,000.

Mash (v2.3; (9)) estimates were gathered using the subcommand `sketch` with the minimum *k*-mer copy number for filtering set to 10 and a `sketch` size of 100,000.

The parameters used for LRGE and Mash were selected based on a sweep of parameter combinations on a validation set described in the appendix (Parameter exploration).

To generate GenomeScope2 estimates, we computed the histogram of *k*-mer frequencies using KMC (v3.2; (13)) with a *k*-mer size of 21, a minimum *k*-mer count (`-ci`) of 2, and a maximum *k*-mer count (`-cs`) of 10,000. These parameters were selected as they are the default recommended in the GenomeScope2 repository. We then ran GenomeScope2 (v2.0.1; (4)) against the frequency histogram with the same settings, with ploidy of 1.

Raven is a genome assembly tool that is designed to be time- and memory-efficient and uses an overlap-layout-consensus approach (12) - keeping with the overlap theme. We set Raven (v1.8.3) to perform no polishing (`-p` 0) and use the size of the computed assembly as the “estimate”.

We evaluated the accuracy of each method using relative error, ϵ_rel_, which measures the percentage difference between the estimated value (*Ĝ*) and the true value (*G*), scaled relative to the true value. It quantifies how close the estimate is to the true value, with positive values indicating overestimation and negative values indicating underestimation. It is defined as

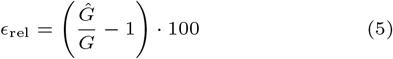

Statistical testing of the absolute relative error, CPU time, and maximum memory usage differences between methods were performed using a pairwise Tukey’s range test. For the absolute relative error testing, we only compared between the same sequencing technology – i.e., we do not compare PacBio for method A with ONT for method B.

#### Prokaryote evaluation

We performed a large-scale evaluation of the estimation methods using a set of 3370 publicly-available bacterial long-read sequencing runs – 2468 ONT and 902 PacBio. These runs were selected as they were associated with a high-quality RefSeq assembly, which we took to be the true size. A detailed description of the data collection and filtering performed is provided in the appendix (Dataset selection).

#### Eukaryote evaluation

While the primary development focus for LRGE was bacterial genomes, we investigated how the method generalises to eukaryotes, such as multichromosome and diploid organisms. We used ONT reads from three model organisms with a high-quality RefSeq genome: *Saccharomyces cerevisiae, Drosophila melanogaster*, and *Arabidopsis thaliana*.

Due to the larger genome sizes of these organisms, we used 10,000 query reads and 20,000 target reads for the *2set* strategy on *S. cerevisiae* and default (25,000) for the *ava* strategy. For *D. melanogaster* and *A. thaliana*, we used 100,000 target reads and 50,000 query reads for the LRGE *2set* approach and 100,000 reads for the *ava* strategy. The parameters were kept the same as for the prokaryote evaluation for all other methods.

## Results

To evaluate the accuracy of the estimations produced by LRGE, we collected 3370 publicly-available bacterial long-read sequencing runs with associated high-quality reference assemblies (see Prokaryote evaluation). We benchmarked the two LRGE strategies against three other methods: Mash, which uses sketching of *k*-mers to produce an estimate; GenomeScope2, which performs statistical analyses on *k*-mer frequency spectra; and Raven, which is a rapid genome assembly method (see Evaluation.

Our primary metric of comparison is relative error (ϵ_rel_; Equation 5), which measures the percentage difference between the estimate and the true value, scaled by the true value. For example, a ϵ_rel_ of 50 for a sample with a genome size of 2 Mbp indicates the estimate is over by half the true size – 3 Mbp. To simplify the evaluation, we use the absolute value of ϵ_rel_ to make for easier comparison across samples.

Figure 1 shows the results of this comparison. The first observation is that the two LRGE strategies provide very similar estimates, with no significant difference. Second, LRGE and Raven perform better with ONT data than PacBio, while Mash and GenomeScope2 are the opposite. Raven provides the best genome size estimates, being statistically significantly better than all methods on PacBio data (mean: 3.5%, median: 2.6%), and GenomeScope2 and Mash on ONT data. (mean: 2.9%, median: 1.1%) LRGE had significantly lower ϵ_rel_ compared to GenomeScope2 and Mash for ONT data (*ava* : mean 9.2%, median 4.6%; *2set* : mean 8.9%, median 4.8%), though the inverse was true for PacBio data. The full results can be seen in Supplementary Table S1.

**Fig 1.**
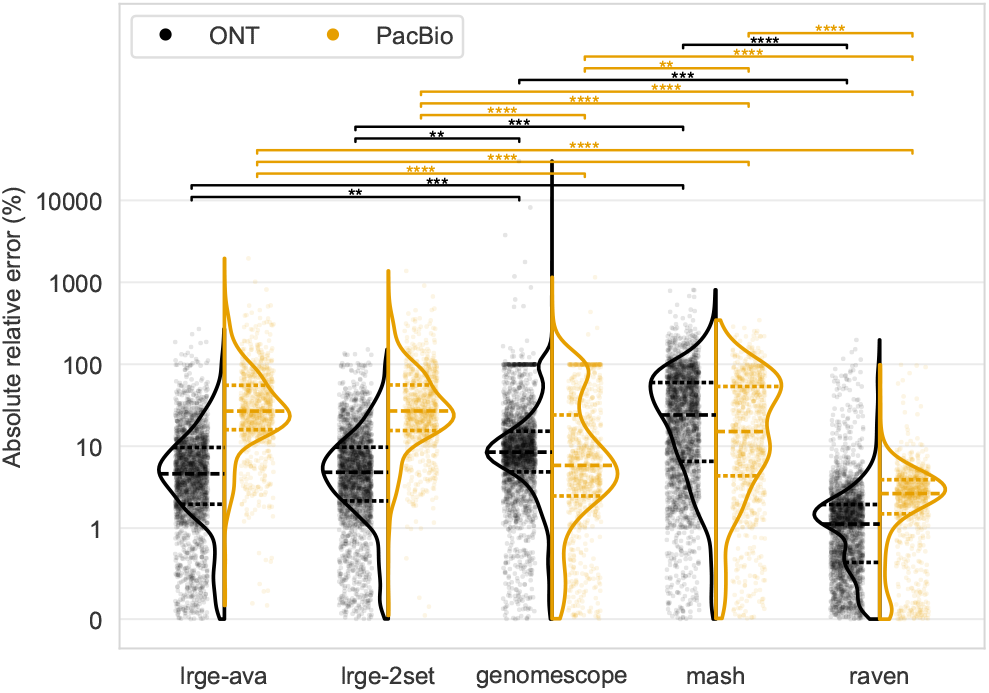
Absolute relative error (y-axis) for each method’s (x-axis) genome size estimation on ONT (black) and PacBio (orange) data. The y-axis is scaled according to a symmetric logarithm, which is linear between −1 and 1 and logarithmic (base 10) thereafter. The statistical annotations are the result of a Tukey’s range test and are coloured by the sequencing platform being compared. The dashed lines in the violins are the quartiles.

### Outliers

GenomeScope2 and Raven tended to underestimate the genome size (Figure S4) on ONT data, though the median was approximately −1% for Raven. In terms of outliers for LRGE there were two common patterns. Major underestimates were found to be cases where there was dramatically disproportionate depth across the genomes. In one example, a *Pandoraea fibrosis* run (SRR9733840), the ribosomal RNA genes had up to 100,000x read depth. In another, a *Enterobacter ludwigii* run (SRR12247681) had two plasmids with 50,000x and 160,000x depth. This dramatic difference in copy number for plasmids is a known mechanism of transient antibiotic heteroresistance (14). These dramatic depth differences lead to underestimates because, when randomly selecting reads, there is a high likelihood that mostly plasmid or gene reads will be selected, and thus the estimate will reflect the (much smaller) size of the plasmid(s) or gene(s).

The other major outlier pattern was low quality runs leading to large overestimates. This is illustrated in Figure S5 where we see runs with a ϵ_rel_ over 50 with average read qualities much lower than other samples. Assumably this lower quality makes it very difficult to overlaps reads, leading to fewer overlaps, and thus larger genome size estimates.

### Calibration of estimate confidence range

As we calculate a genome size estimate for each read, we can determine a range within which the estimate is likely to fall with a given level of confidence. To achieve this, we scanned all percentile ranges of width 50%, ensuring that the overall estimate (median) was contained within the range. Figure S6 illustrates this process and shows that the percentile range that maximizes the proportion of samples with the true genome size within the corresponding range is 15–65% for both LRGE strategies. We can therefore say that we are 92.5% (*2set*) and 87.8% confident the genome size lies within this estimate range. On the prokaryote dataset, this range gave an interquantile range (IQR) of 0.53 and 0.44 relative size 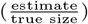 for the *2set* and *ava* strategies, respectively. For example, an IQR of 0.53 means that for the *2set* strategy, the middle 50% of estimates fall within a range that is 53% of the true genome size, while for the *ava* strategy, this range is only 44%. This illustrates that the *ava* strategy provides a more precise estimate, with less variability in the middle half of the predictions compared to the *2set* strategy.

### Runtime and memory benchmark

Figure 2 shows the benchmark of CPU time and maximum memory usage for all methods. In terms of CPU time, LRGE *2set* (mean: 38.2s, median: 17.4s) and GenomeScope2 (mean: 31.9s, median: 31.6s) had the quickest runtime, with no statistically significant difference between the two. Raven was significantly slower than all methods except LRGE *ava*. The longest recorded CPU time, 2.4 hours, was LRGE *ava*. Both LRGE strategies had large CPU time outliers, highlighted by the differences between their mean (*ava* : 194.7s, *2set* : 38.2s) and median (*ava* : 77.2s, *2set* : 17.4s). These outliers were enriched in samples with large underestimates (Figure S7). Together with the fact that samples with large overestimates tended to have very fast runtimes (Figure S7), this indicates that the runtime is proportional to the number of overlaps (see Outliers).

**Fig 2.**
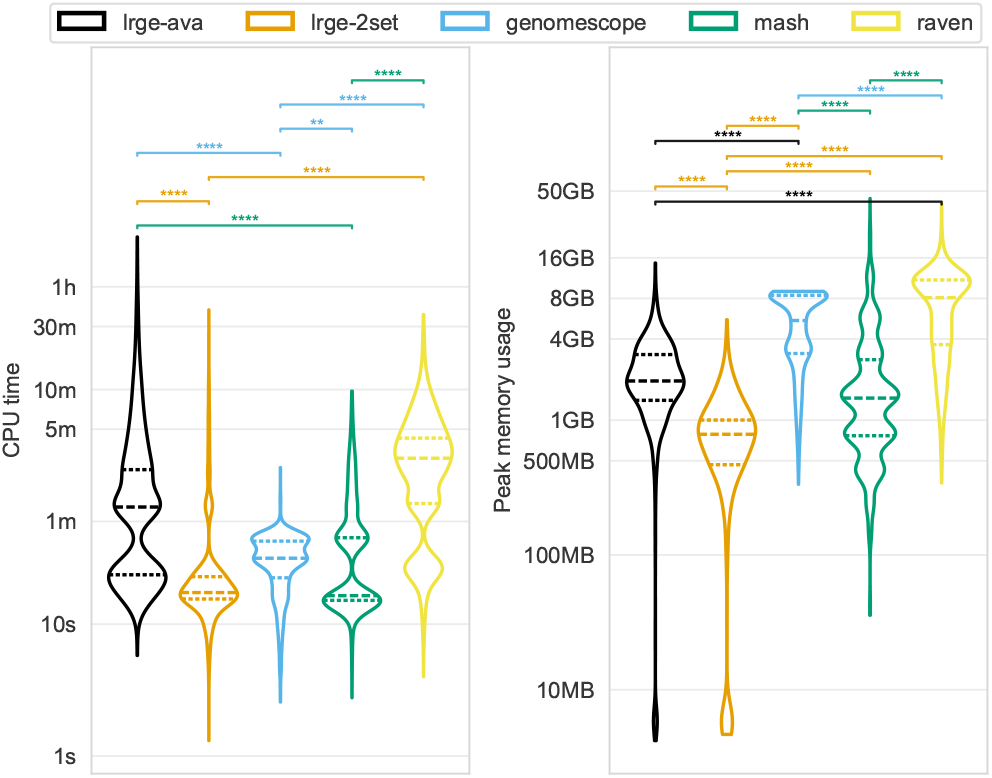
CPU time (left y-axis) and maximum memory usage (right y-axis) for each method (colours). The y-axis is log-scaled (base 10). The statistical annotations are the result of a Tukey’s range test and are coloured by the method with the lower mean value. The dashed lines in the violins are the quartiles.

LRGE *2set* memory usage (mean: 776 MB, median: 749 MB) was significantly lower than all other methods, while Raven had the highest memory usage (mean: 7.2 GB, median: 7.7 GB). The highest recorded memory usage was Mash, with a maximum of 42 GB.

The full results can be seen in Suppl. Table S2.

### Eukaryotes

While the primary focus and development of this work has been bacteria-focused, we investigated how LRGE generalises to multi-chromosome and diploid organisms. We evaluated LRGE, along with the other three methods, against three model organisms: *S. cerevisiae, D. melanogaster*, and *A. thaliana* (see Eukaryote evaluation). These results are presented in Table 1 and show that LRGE indeed generalises well to eukaryotic organisms, having the lowest ϵ_rel_ for *S. cerevisiae* and *A. thaliana*, and a ϵ_rel_ within 3-8% on *D. melanogaster*. The full results can be seen in Suppl. Table S3.

**Table 1.** Estimates on eukaryote samples.

| Organism | Ploidy | Chromosomes | True size* | Method | Estimate* | $\epsilon_{\text{rel}}^{\dagger}$ | Time<sup>‡</sup> | Memory (GB) |
| --- | --- | --- | --- | --- | --- | --- | --- | --- |
| *Saccharomyces cerevisiae* | 1 | 16 | 12.2 Mbp | LRGE *2set* | 12.0 Mbp | **-0.97** | 0:07:26 | 4.3 |
|  |  |  |  | LRGE *ava* | 12.0 Mbp | -1.24 | 0:11:18 | 5.2 |
|  |  |  |  | Mash | 55.7 Mbp | 358.48 | **0:05:04** | **0.9** |
|  |  |  |  | GenomeScope2 | 13.7 Mbp | 12.89 | 0:10:30 | 11.2 |
|  |  |  |  | Raven | 13.4 Mbp | 10.19 | 19:26:54 | 36.7 |
| *Drosophila melanogaster* | 2 | 7 | 143.7 Mbp | LRGE *2set* | 138.9 Mbp | -3.33 | 0:14:45 | 7.6 |
|  |  |  |  | LRGE *ava* | 132.6 Mbp | -7.77 | 0:17:19 | 8.5 |
|  |  |  |  | Mash | 141.4 Mbp | **-1.62** | **0:10:09** | **1.8** |
|  |  |  |  | GenomeScope2 | 128.3 Mbp | -10.71 | 0:18:12 | 11.2 |
|  |  |  |  | Raven | 135.0 Mbp | -6.07 | 37:08:46 | 61.8 |
| *Arabidopsis thaliana* | 2 | 5 | 119.7 Mbp | LRGE *2set* | 143.4 Mbp | 19.80 | 0:19:11 | 8.6 |
|  |  |  |  | LRGE *ava* | 142.9 Mbp | **19.43** | 0:19:56 | 9.3 |
|  |  |  |  | Mash | 245.6 Mbp | 105.27 | **0:16:00** | **1.8** |
|  |  |  |  | GenomeScope2 | 162.7 Mbp | 35.98 | 0:31:11 | 11.3 |
|  |  |  |  | Raven | 192.3 Mbp | 60.74 | 74:33:40 | 94.2 |

\* Size values have been rounded to megabase pair with one decimal place for compactness.
<sup>†</sup> $\epsilon_{\text{rel}}$ is relative error expressed as a percentage, as defined in Equation 5.
<sup>‡</sup> CPU time formatted as [h]:mm:ss.
**Bold text** indicates the best value for the relevant metric for each organism.

## Conclusion

LRGE provides a robust and efficient method for genome size estimation tailored to long-read sequencing technologies. By leveraging read-to-read overlap data, it circumvents the limitations of *k*-mer-based methods and performs well across diverse datasets, including bacteria and eukaryotes. LRGE outperforms existing *k*-mer-based tools in both accuracy and computational efficiency and delivers estimates comparable in accuracy to those derived from an assembly-based approach, Raven, but with significantly reduced computational resource requirements. Its flexibility and scalability make LRGE a valuable addition to the bioinformatics toolkit, supporting applications in genome assembly, evolutionary studies, and sequencing depth estimation.

## Competing interests

No competing interest is declared.

## Author contributions statement

**Michael B Hall:** Conceptualization, Data curation, Formal Analysis, Investigation, Methodology, Software, Writing – original draft, Visualization, Writing – review & editing. **Lachlan J M Coin:** Conceptualization, Formal Analysis, Funding acquisition, Investigation, Methodology, Software, Supervision, Writing – original draft, Writing – review & editing.

## Acknowledgment

This research was supported by the University of Melbourne’s Research Computing Services and the Petascale Campus Initiative. We thank Ryan Wick for insightful suggestions and Leah Roberts for discussions relating to large plasmid copy numbers.

## Funding

This work was supported, in part, by the Australian Government Medical Research Future Fund (MRFF) Genomics Health Futures Mission (GHFM) Flagships – Pathogen Genomics Grant (FSPGN000045) META-GP: DELIVERING A

### CLINICAL METAGENOMICS PLATFORM FOR AUSTRALIA

The funding body had no role in the design, analysis, interpretation, or writing of this work.

## Data availability

All code and metadata required to perform the analyses in this work are available on GitHub at https://github.com/mbhall88/lrge and are archived at Zenodo (15) or in Supplementary Table S3 for the eukaryote samples.

## Parameter exploration

We explored the best combination of parameters to be used for the larger dataset in the main text, by validating on a much smaller dataset. For this, we used ONT reads from 14 different bacterial species (16). In particular, these are R10.4.1 simplex reads basecalled with the v4.3.0 Dorado super accurate (sup) model (17).

### LRGE two-set overlaps

For the LRGE two-set (*2set*) approach, the three parameters we explore are the number of query and target reads used for the overlaps, *Q* and *T*, respectively, and the read selection strategy.

For the *long* strategy, we select the |*Q*| longest reads, *Q* and the next |*T*| longest reads, *T*. In the *rand* strategy, we randomly select *Q* and *T* reads, ensuring these sets are disjoint, using Rasusa (v2.1.0) (10). We then determine the overlaps using minimap2 (v2.28) (11) with the *Q* reads as the query and *T* as the target. For the *rand* strategy, we performed three replicates for each sample-*Q*-*T* combination.

The metrics we used to assess the performance of the different values of *Q* and *T* are relative error (Equation 5; ϵ_rel_) and the coefficient of determination (*R*^2^), with respect to the identity line, for each parameter combination. From Figure S1 we can see clearly that the *long* strategy is inferior to the *rand* strategy. *Q* = 5000 with *T* = 10000 had the best balance of *R*^2^ and ϵ_rel_ and were therefore selected as the default.

### LRGE all-vs-all overlaps

For the LRGE all-vs-all (*ava*) approach, the two parameters we explore are the number of reads used for the overlaps, *n*, and the read selection strategy. For the *long* strategy, we select the *n* longest reads and in the *rand* strategy, we select *n* reads at random using Rasusa (v2.1.0) (10). We then determine the overlaps using minimap2 (v2.28) (11) with the *n* reads as both the target and query. For the *rand* strategy, we performed three replicates for each sample-*n* combination.

From Figure S2 we can see clearly that the *long* strategy is inferior to the *rand* strategy. *n* = 10000 had the highest *R*^2^ and closest ϵ_rel_ to zero, though *n* = 25000 was very close. We chose to use 25000 as the default because of the fact it was so close to *n* = 10000 and the fact that it performed much better on the *long* strategy.

### Mash

The parameters we explored for Mash (9) were the sketch size (`-s`) and the minimum copies of a *k*-mer required to pass the noise filter (`-m`). We ran the Mash subcommand `sketch` with these parameters for each of the 14 samples. Figure S3 shows the results of this parameter exploration, with a sketch size of 100000 and minimum copies of 10 giving the tightest distribution of relative error near zero.

## Dataset selection

Metadata for all complete bacterial assemblies in NCBI’s RefSeq (excluding MAGs) were downloaded using NCBI’s `datasets` command line tool. We selected assemblies where: (i) the sequencing technology section mentions PacBio or Oxford Nanopore Technologies, (ii) the assembly release date was after 01/01/2016, (iii) CheckM contamination was below 3%, (iv) the CheckM completeness was greater than 95%, (v) the CheckM completeness percentile for the species was above 75%. Next, we gathered metadata on the sequencing runs associated with the BioSample accession for each assembly. We removed assemblies which did not actually have any long read ONT or PacBio sequencing, despite criteria (i) from above.

The long read data left after these filtering steps were downloaded using Kingfisher (v0.4; (18)). We then removed a further 1,130 sequencing runs due to low volume of sequencing reads, indicated by the readset not having sufficient reads to run LRGE *2set*, or GenomeScope2 giving an error stating it was unable to converge on an estimate (an error that typically indicates low sequencing depth). We also removed 22 runs which were actually Illumina reads, despite being labeled as PacBio or ONT on the NCBI SRA, and 16 runs for which a valid FTP address could not be found to download the data.

In the end, we were left with 3,370 long read sequencing runs.

---

**Fig. S1.**
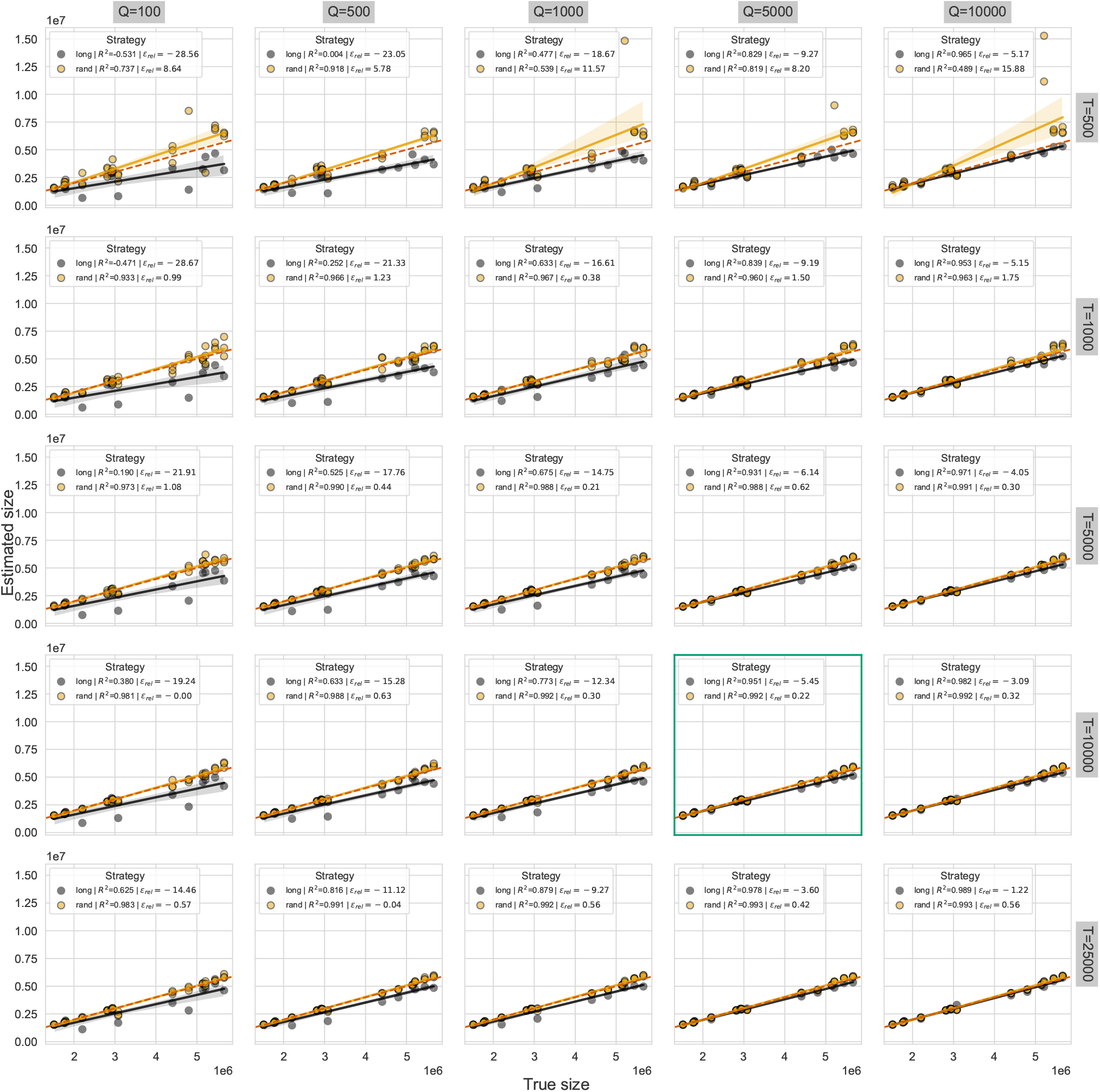
LRGE two-set parameter exploration. Columns are the query read set size *Q*, while the rows are the target read set size *T*. Each subplot shows the estimated genome size (y-axis) against the true genome size (x-axis), with points coloured by the strategy. ϵ_rel_ is the relative error and *R*^2^ is the coefficient of determination. The subplot with the green border is the one that was selected as the default parameters.

**Fig. S2.**
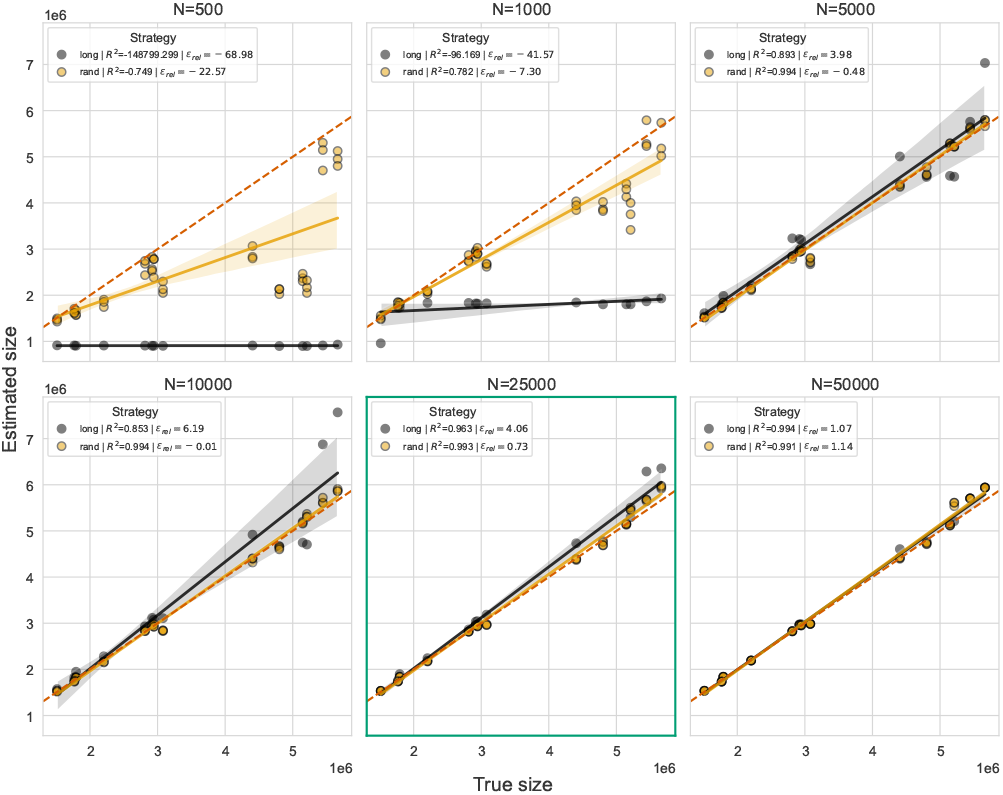
LRGE all-vs-all parameter exploration. Each subplot represents the number of reads (*N*) used and shows the estimated genome size (y-axis) against the true genome size (x-axis), with points coloured by the strategy. ϵ_rel_ is the relative error and *R*^2^ is the coefficient of determination. The subplot with the green border is the one that was selected as the default parameters.

**Fig. S3.**
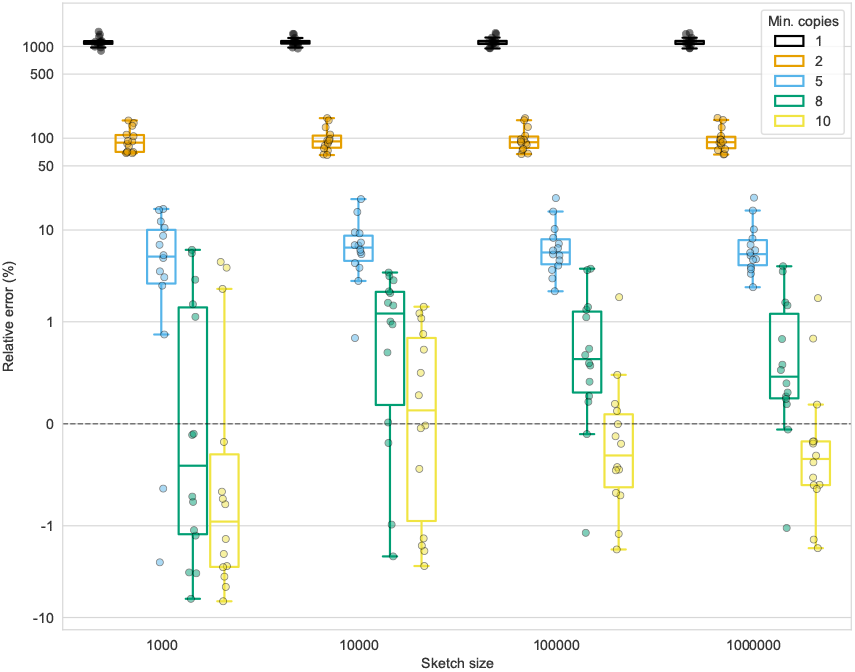
Exploring the best-performing combination of Mash sketch size (x-axis) and minimum required copies of a *k*-mer to pass noise filtering (colours). The darker horizontal dashed line at *y* = 0 indicates the optimal relative error. The y-axis is scaled according to a symmetric logarithm, which is linear between −1 and 1 and logarithmic (base 10) thereafter.

**Fig. S4.**
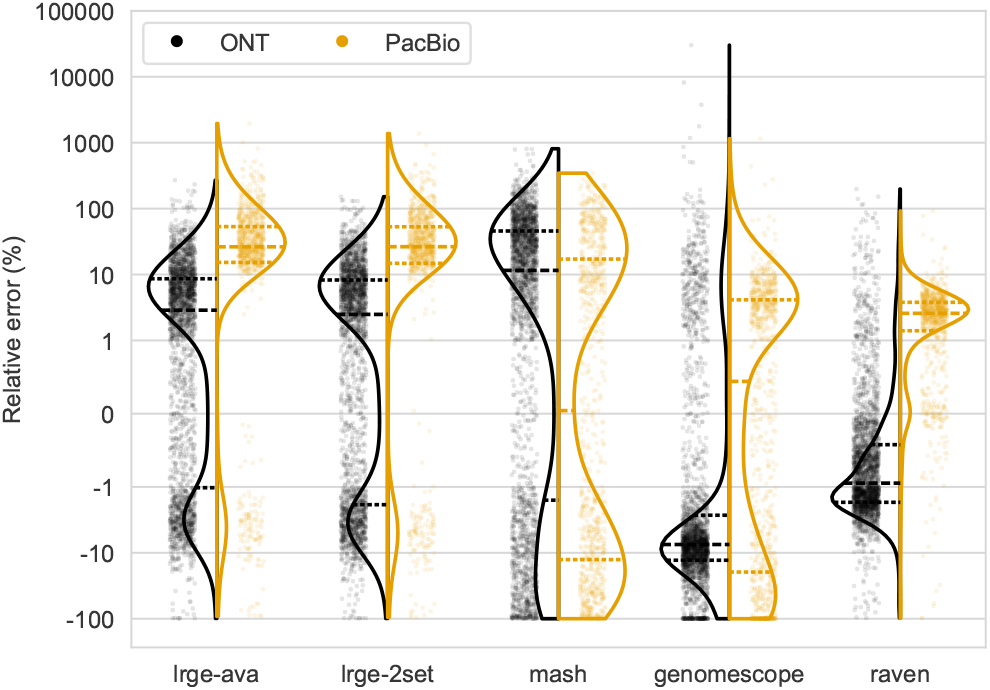
Relative error (y-axis) for each method’s (x-axis) genome size estimation on ONT (black) and PacBio (orange) data. The y-axis is scaled according to a symmetric logarithm, which is linear between −1 and 1 and logarithmic (base 10) thereafter. The dashed lines in the violins are the quartiles.

**Fig. S5.**
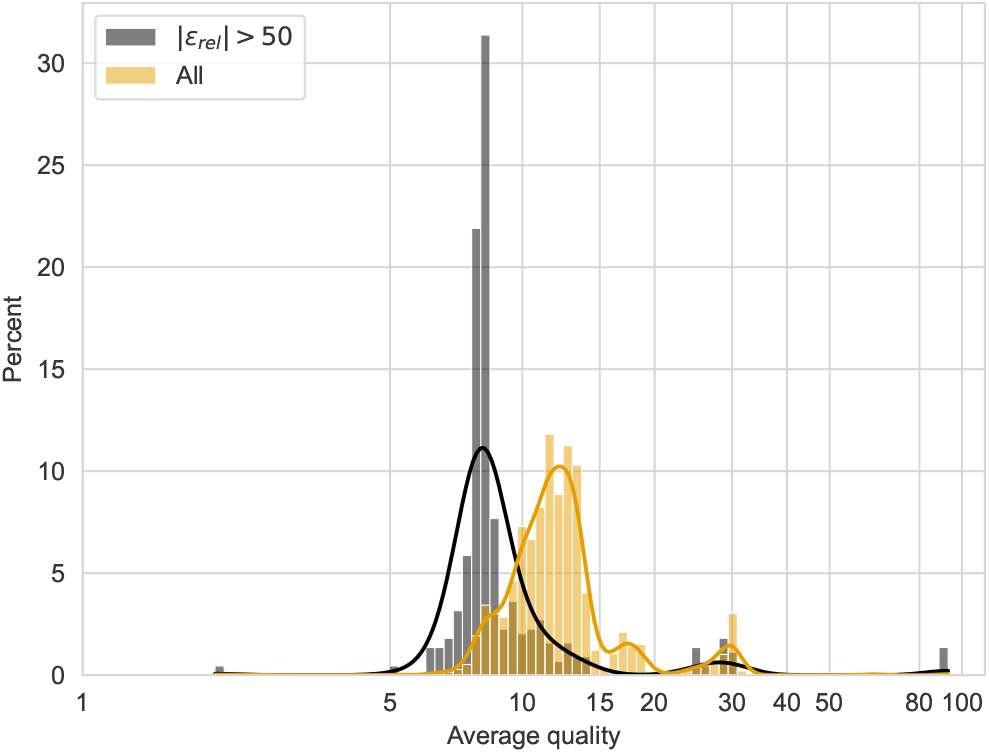
Distribution of average read quality for samples with a relative error (ϵ_rel_) greater than 50 (black) and all other samples (orange).

**Fig. S6.**
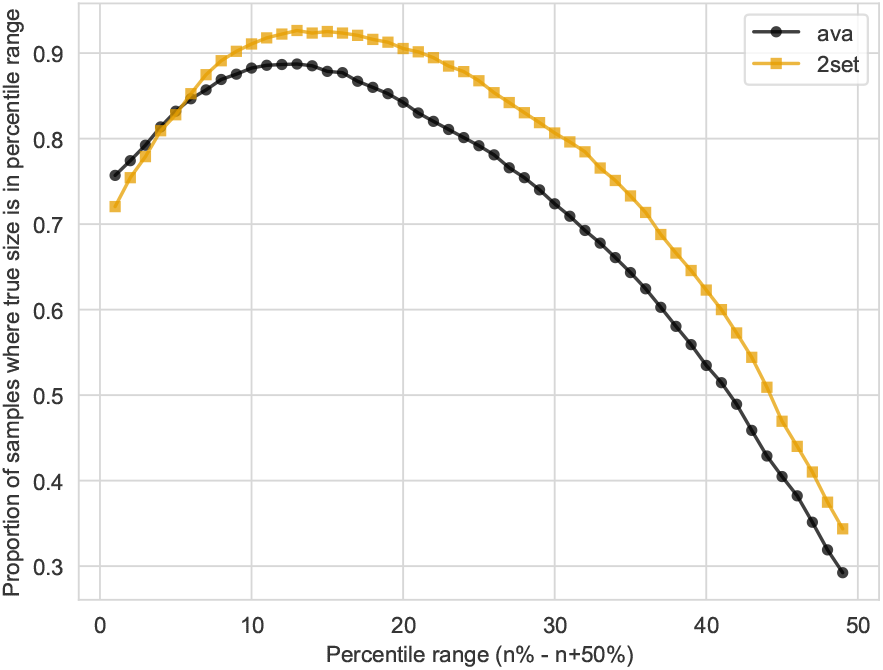
Calibrating the optimal percentile range for LRGE estimates. The x-axis represents the lower end of the range. So an x value of 10 means a percentile range of 10-60. The y-axis represents the proportion of samples for which the true genome size lies within the corresponding percentile range on the x-axis. The two colours represent the two LRGE strategies.

**Fig. S7.**
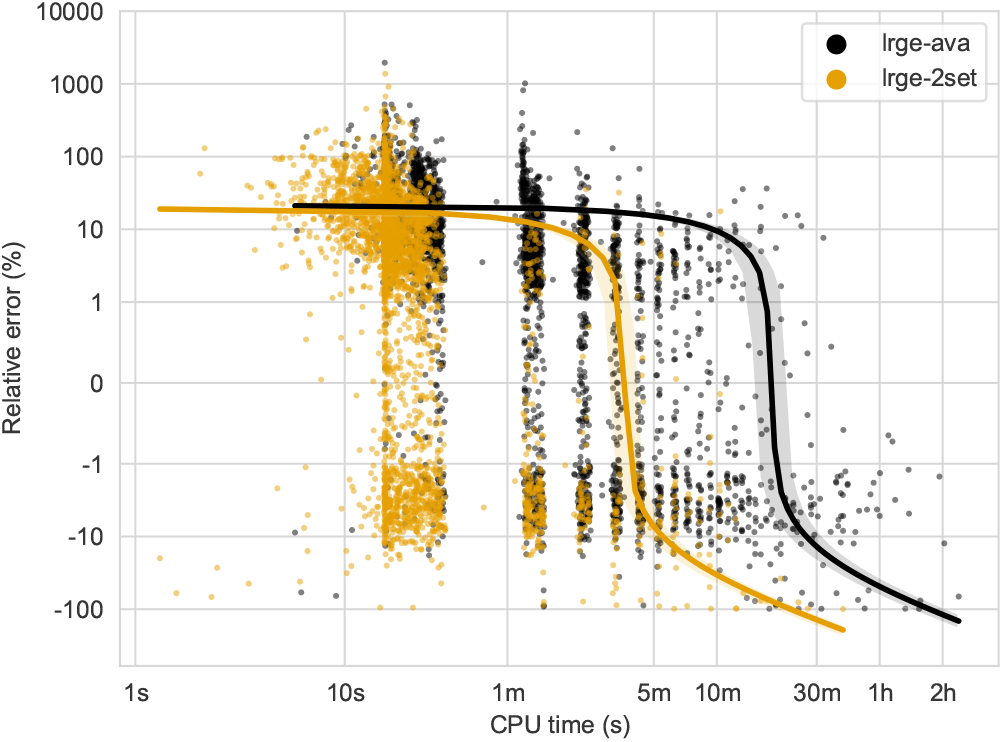
Relationship between CPU time (x-axis) and relaive error (y-axis) for LRGE strategies (colours). The line represents a linear regression model fit to the data.

## Notes

https://github.com/mbhall88/lrge

### Competing Interest Statement

The authors have declared no competing interest.

